# Improving leukemic CD34^+^/CD38^−^ blasts characterization with single-cell transcriptome sequencing

**DOI:** 10.1101/141754

**Authors:** Ambra Sartori, Phil Cheng, Emilie Falconnet, Pascale Ribaux, Jean-Pierre Aubry-Lachainaye, Mitchell P. Levesque, Stylianos E. Antonarakis, Thomas Matthes, Christelle Borel

## Abstract

Acute myeloid leukemia (AML) is a particularly aggressive blood cancer that is difficult to treat because of the incomplete eradication of rare blast cells that possess self-renewal and leukemia-initiating properties. To characterize resistant blasts, we analyzed for the first time the transcriptomes of individual CD3^4+^/CD38^−^ blasts by single-cell mRNA sequencing of 359 CD33^+^/CD34^+^/CD38^−/+^ sorted cells from two patients with AML and four unaffected individuals. We demonstrated that the captured blasts possess the transcriptomic hallmarks of self-renewal and leukemia-initiating ability. The effects of somatic mutations on the cancer cells are visible at the transcriptional level, and the cellular signaling pathway activity of the blasts is altered, revealing disease-associated gene networks. We also identified a core set of transcription factors that were co-activated in blasts, which suggests a joint transcription program among blasts. Finally, we revealed that leukemogenesis and putative prognostic gene-expression signatures are present at diagnosis in leukemic CD33^+^/CD3^4+^/CD38^−^ cells and can be detected using a single-cell RNA sequencing approach.

## BACKGROUND/ INTRODUCTION

Acute myeloid leukemia (AML) is a particularly aggressive blood cancer that affects blood cell precursors generated in bone marrow [1, 2]. In 1997, D. Bonnet and J. E. Dick observed that a CD34^+^/CD38^−^ blast cell population was able to reconstitute human acute leukemia in non-obese diabetic/severe combined immunodeficiency mice (NOD/SCID) [3, 4]. Thus, the authors referred to these blast cells as leukemic stem cells (LSCs) based on their analogous relationship to the normal self-renewing hematopoietic stem cells (HSCs) that reconstitute normal hematopoiesis [3, 5–7]. LSCs are rare (< 10% of malignant CD34^+^ leukemic blast cells) [8] and appear hidden among the heterogeneous bulk of the tumor cell mass; in addition, they are capable of self-renewal and partial differentiation but remain restricted to an immature phenotype similar to normal HSCs. Despite intensive research efforts focused on the precise characterization of the phenotype and functional capabilities of LSCs, these cells have remained somewhat elusive because they are rare and impossible to analyze with standard genomic procedures that require at least 100’000 cells [9–11]. This is exemplified by conflicting results on the precise phenotype of LSCs and their absence of cell surface marker [12]. Nevertheless, current models associate an abundance of LSCs with refractory disease and poor clinical outcome, which supports the importance of understanding the phenotype and functional properties of CD33^+^/CD34^+^/CD38^−^ blasts [11–14].

With the advent of next-generation sequencing and the improvement of single-cell sorting methods from body fluids and tumor tissues, the characterization of the transcriptomes of rare cells has become possible [15] [16][17, 18]. We analyzed a total of 359 CD33^+^/CD34^+^/CD38^−/+^ single cells from two patients with AML and four patients with normal CD33^+^/CD34^+^/CD38^−/+^ single cells. We obtained results that would be impossible to achieve using a standard bulk RNA-seq technology because of a suboptimal statistical power due to a limited sample size of only six individuals including 4 non-AML. The transcriptomic analysis of hundreds of cells enables to reveal a co-expressed set of genes and signaling pathways unique to each of the two patients with AML. Furthermore, these data provide insight into the disease state and molecular biology of AML.

## RESULTS

The goal of this study was to examine the transcriptional landscape of CD34^+^/CD38^−^ blasts by comparing them to CD34^+^/CD38^+^ blasts and to normal CD34^+^/CD38^−^ cells. We used flow cytometry to sort CD33^+^/CD34^+^/CD38^−^ and CD33^+^/CD34^+^/CD38^+^ blast populations from the bone marrow cell suspensions of two patients with AML (AML1 (M0) and AML2 (M5)) and four patients with normal bone marrow (N) (refer to the Materials and Methods), and we then performed single-cell mRNA sequencing on 359 sorted cells. After stringent filtering, we generated 311 single-cell RNA-seq profiles with an average of 7 × 10^6^ uniquely mapped reads per cell (Figure 1A and B, see the Materials and Methods). As expected, the number of detected genes per cell was comparable between the conditions and variable between the cells [19, 20] (Figure 1). On average, 1764 transcribed genes were detected per cell (RPKM > 10). Consistent with previous reports [19–21], we observed a substantial variability in the cell-to-cell transcriptome (Pearson 0.0007 < r < 1, mean 0.57) (Figure 1C). Bulk RNA from each individual was prepared for bulk-to-single cell transcriptome correlations. As described, the bulk transcriptomes were highly correlated with the single-cell transcriptomes (Pearson 0.38 < r < 0.89, mean 0.63) (Figure 1C), thus supporting the assumption that bulk samples reflect the average of single-cell populations [20, 22].

**Figure 1.**
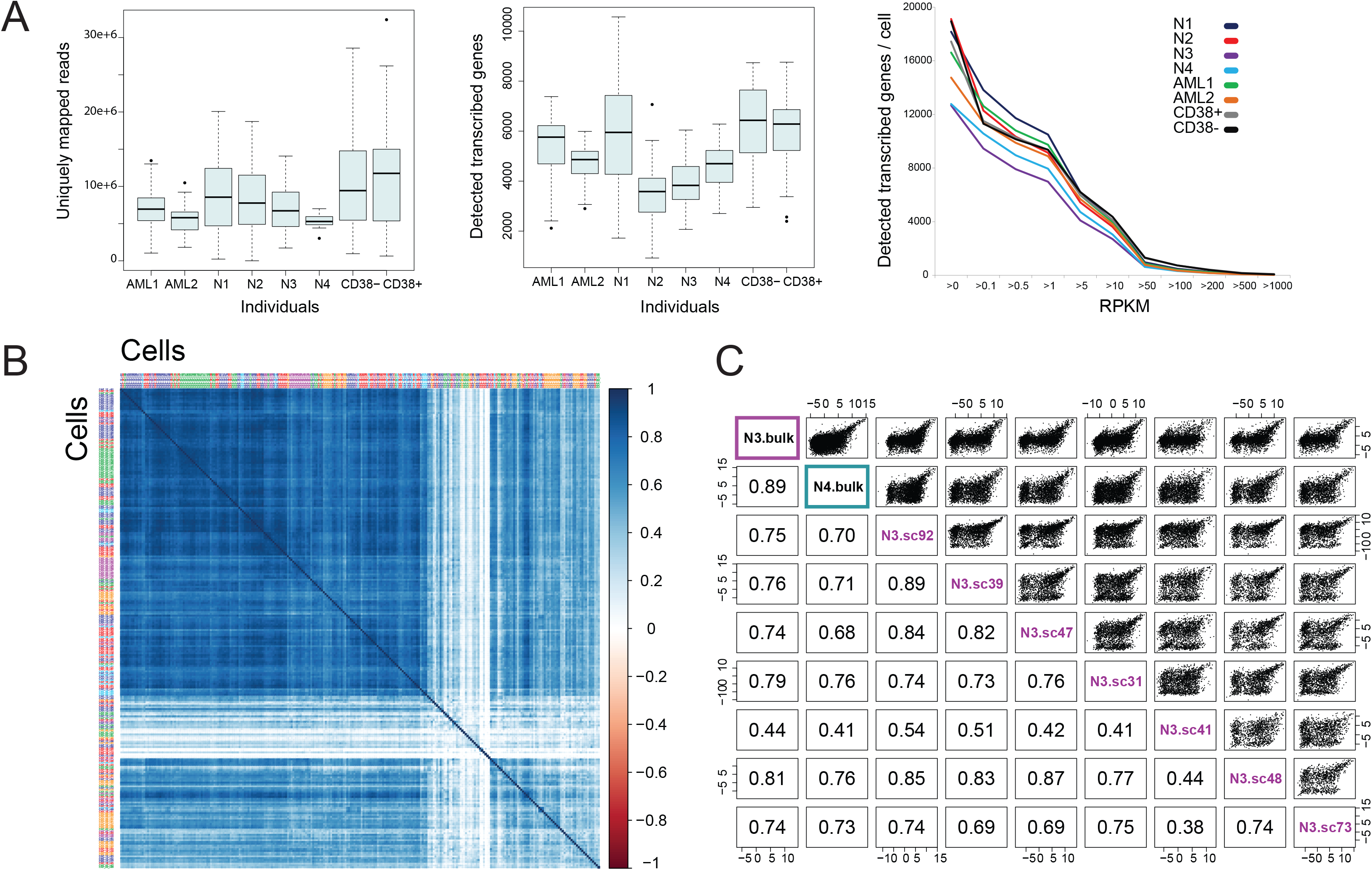
Heterogeneity in single-cell measurements of transcript expression. **(A)** Boxplots showing the number of uniquely mapped reads and transcribed genes detected in 313 single cells from 4 healthy patients (N1, N2, N3, N4) and 2 patients with AML (AML1, AML2). CD38^−^ and CD38^+^ blast cells from patient AML2 were sorted separately (refer to the Materials and Methods). See also Figure S1. The right panel shows the cumulative average number of transcribed genes detected per cell for each sample and per RPKM category. **(B)** Hierarchical clustering of 267 single-cell samples (refer to the Materials and Methods) based on Pearson correlation. The cell labels are colored according to the sample origin (same color code as the right panel in (A)). CD38^−^ and CD38^+^ single cells are not included in this plot. The correlation coefficient is also colored according to the scale ranging from 1.0 (blue) to - 1.0 (red). **(C)** Cell-to-cell and cell-to-bulk correlation matrix including 7 single-cell samples (sc) from N3 and bulk samples from N3 and N4. Scatter plots show correlation with gene expression (RPKM > 0). Numbers represent pairwise Pearson’s correlation coefficients.

### Individual leukemic CD34^+^/CD38^−^ cells trigger pathways that promote stemness and cancer progression

We first compared the transcriptome profiles of 24 CD34^+^/CD38^−^ cells and 24 CD34^+^/CD38^+^ cells from the same patient with AML (AML2), and an analysis of the differentially expressed transcripts identified 625 genes that presented significant changes (p-value < 0.05) (refer to the supplementary data, Table S1). As expected, the hierarchical clustering map of the differentially expressed transcripts showed two distinct cell populations (Figure 2A). An analysis of the gene ontology terms conducted on this set of differentially expressed transcripts revealed significantly enriched terms associated with the genes mainly implicated in the NOD-like receptor pathway (e.g., *CXCL2, CXCL8, NLRP3*, and *TRAF6*) (Figure 2B). This term also includes chemokines that are critical for the survival and proliferation of cancer cells, such as CXC-motif ligand 8 (*CXCL8, IL-8*) [23, 24]. Malignant CD34^+^/CD38^−^ cells displayed reduced cell cycle activity, and genes that promote cell proliferation and cell cycling (*CDK7, CDKN2A, HDAC1, MCM3, PCNA*, and *MYC*) were downregulated compared with the more differentiated blasts. Interestingly, DNA replication genes (*POLA2, RFC3*, and *RNASEH2A*) were activated in those cells, whereas the nucleotide excision repair pathway was significantly downregulated, thus suggesting a promoted DNA damage (Figure 2B), which was previously described in quiescent cells [25–27]. A gene set enrichment analysis (GSEA) was performed to evaluate the presence of well-characterized stem cell regulatory genes in our dataset (Figure 2C). We tested predefined gene lists from published gene expression profiles of pathways activated in LSCs and HSCs (refer to the supplementary data, Table S4) [11, 12, 28, 29] and confirmed that the stem-associated genes expressed by our sorted CD33^+^/CD34^+^/CD38^−^ leukemic cells were consistent with published data (FDR < 0.01). Several studies have implicated the Wnt/β-catenin pathway in the development of leukemia stem cells [30]. A GSEA of this specific pathway confirmed the overexpression of 49 of 148 genes in CD34^+^/CD38^−^ blasts (Figure 2C).

**Figure 2.**
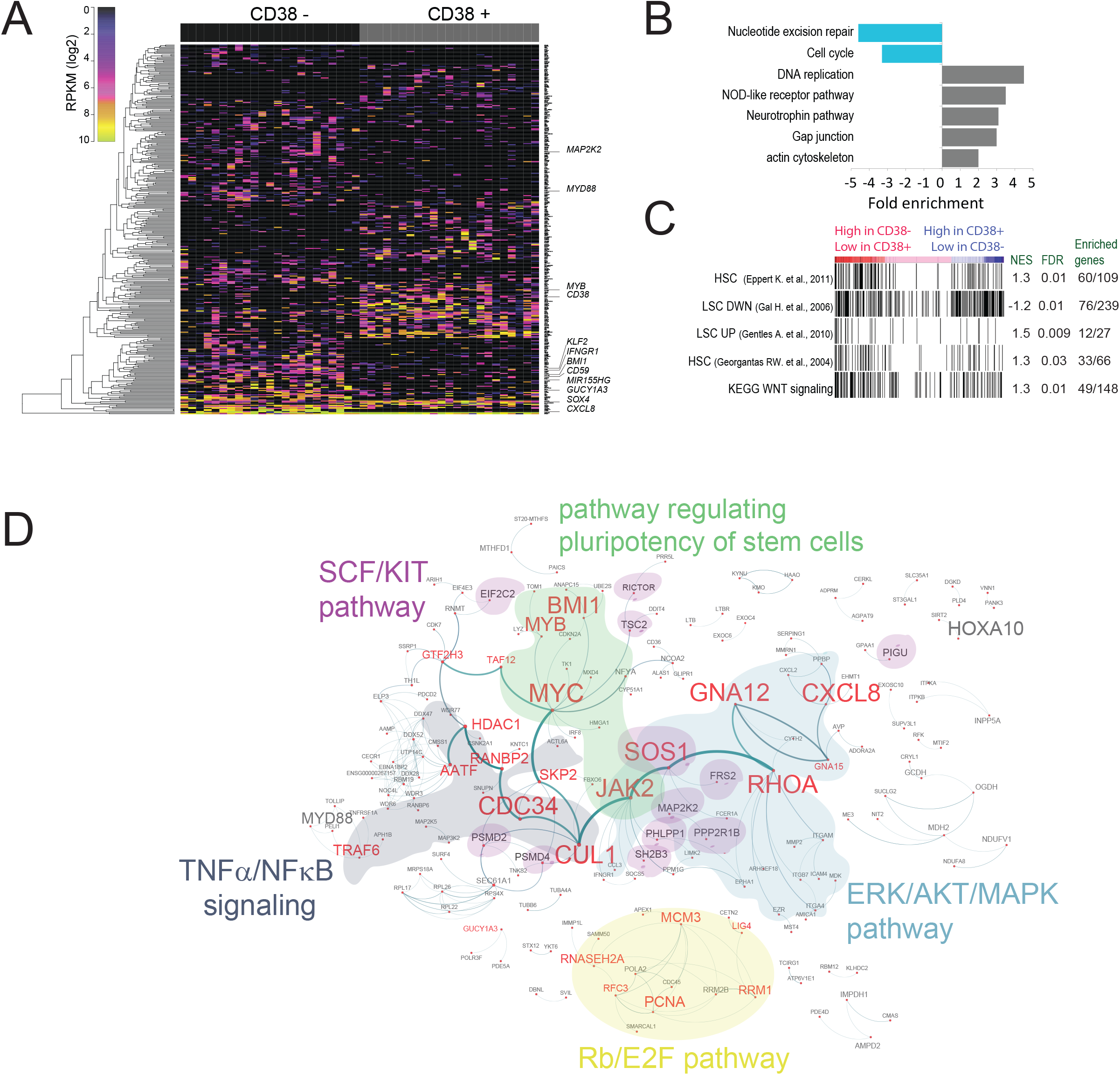
Leukemic CD34^+^/CD38^−^ blasts display a stemness signature when compared with leukemic CD34^+^/CD38^+^ blasts from the same AML patient. **(A)** Heat map and supervised hierarchical clustering of differentially expressed genes between CD38^−^ and CD38^+^ cells (refer to the supplementary data, Table S1 for gene list details). The rows represent genes and columns represent single cells from CD38^−^ and CD38^+^ subpopulations. Color coding denotes log2-transformed RPKM values. **(B)** GO term analysis of CD38^−^ and CD38^+^ differentially expressed transcripts (fold enrichment DAVID analysis). The bar diagram shows the significantly enriched terms (p-value < 0.05, Fisher’s Exact Test) among downregulated transcripts (blue) and upregulated transcripts (grey). See also Figure S1. **(C)** Signature enrichment plots from GSEA using 5 different gene target lists (refer to the supplementary data, Table S4 for gene list details). Black vertical bars are genes ordered according to their fold change expression. Values indicate normalized enrichment score (NES), FDR adjusted p-value and number of significant enriched genes out of the total genes tested. **(D)** Gene network visualization of differentially expressed genes between leukemic CD38^−^ and CD38^+^ single cells. Stemness-related genes and pathways are highlighted. See also Figure S2.

The network map illustrates the functional connectivity among the differentially expressed genes (Figure 2D). CD34^+^/CD38^−^ leukemic cells expressed genes related to the following 4 distinct signaling pathways: TNFα/NF-κB, c-Kit–mediated stem cell factor (SCF), Rb/E2F, ERK/MAPK and AKT. Remarkably, genes important for hematopoiesis and leukemogenesis, such as *BMI1* (FC CD38-/CD38+: 4.47, p-value: 0.00113, Kolmogorov-Smirnov statistical test), *HOX* genes (refer to the supplementary data, Figure S2), and *MYB* have been previously reported to be deregulated in leukemic cells [31, 32]. Moreover, these genes have been extensively associated with cancer stem cell maintenance in general [31, 33–35] and support the “stemness” and tumorigenic potential of sorted CD33^+^/CD34^+^/CD38^−^ blasts. Remarkably, expression profiles generated by single-cell RNA-seq on a few number of cells enable to transcriptionally distinguish the molecular features of CD33^+^/CD34^+^/CD38^−^ from CD33^+^/CD34^+^/CD38^+^ blasts.

### CD34^+^/CD38^−^ single-cell transcription profiles distinguish leukemic cells from normal stem cells

To assess whether our framework was suitable for characterizing the transcriptome profiles of CD34^+^/CD38^−^ blasts, we analyzed 267 individual CD33^+^/CD34^+^/CD38^−^ cells from four patients with normal bone marrow and two patients with AML. Interestingly, *SOX4* was among the top 200 highly expressed genes (refer to the supplementary data, Figure S3) in normal CD34^+^/CD38^−^ cells and in the CD34^+^/CD38^−^ blasts of the two AML samples, and its presence was clearly associated with “stemness” properties [36–39], thus reinforcing the suggestion that all of the captured cells might possess the hallmarks of putative stem cells. Remarkably, we identified 5 transcriptionally different clusters on the t-SNE map (Figure 3, Figure S4) (refer to the Materials and Methods). Fifteen cells were unassigned and deliberately allocated to cluster 1. The remaining 219 CD34^+^/CD38^−^ cells grouped into four clusters. AML1 and AML2 cells distinctly clustered in clusters 2 and 4, respectively, whereas all of the non-AML cells formed a distinct and unique cluster, cluster 3 (Figure 3). Cluster 5 consisted of 5 outlier cells from normal patients (Figure 3). The observed clustering indicated that inter-individual variability had less impact on mRNA expression profiles than the disease phenotype and that identified clusters reflect disease states. To identify genes associated with the disease state, the D^3^E method [40] was used to identify differentially expressed transcripts among the AML1, AML2 and non-AML CD34^+^/CD38^−^ cells (Figure 4A). We found 858 and 763 genes that were differentially transcribed in the AML1 and AML2 cells compared with the non-AML cells (p-value < 0.05), respectively, and counted 185 genes that were essentially related to cell cycle regulation and cancer pathways in all cell types (Figure 4B). Among the upregulated or downregulated genes in the AML1 and AML2 cells, the most enriched gene ontology categories were related to the cell cycle, DNA replication, DNA repair, cellular senescence and self-renewal and stemness, such as the JNK pathway, FOXM1 and PLK1 networks, and TGF-beta signaling pathway (Figure 4B). A GSEA performed with literature-based gene lists confirmed that the AML1 and AML2 blasts showed significant enrichment in the published stem cell gene sets and leukemia-activated pathways as well as in a prognostic gene signature (refer to the supplementary data, Figure S5). The overlap of enriched functional-related gene groups in the two AML cells revealed common properties of LSCs in “stemness”-related signaling pathways that control survival, tumorigenesis and self-renewal. The patients with AML1 and AML2 belonged to different subtypes of AML. The leukemic cells of patients with AML2 harbored the common FLT3-ITD mutation, which corresponds to an internal tandem duplication (ITD) in the Fms-like tyrosine kinase 3 gene (*FLT3*), which encodes the receptor for the cytokine FLT3 ligand (FLT3L). Normally, the FLT3 receptor is expressed at the surface of HSCs and is required for the development of myeloid progenitors. The FLT3-ITD mutation results in a hyper-sensitivity of the FLT3 receptor, which promotes uncontrolled cell proliferation mediated by AKT-, MEK-, and ERK-activated pathways [41, 42]. Indeed, we observed that FLT3 transcripts are overexpressed in AML2 cells compared with non-AML and AML1 cells (refer to the supplementary data, Figure S6). The pathways deregulated by this overexpression, i.e., RAS, ERK, AKT, TGF-beta, and GPCR, are illustrated in the networking map (Figure 5A). A comparison with the cellular network map from the AML1 sample showed that these perturbations were specific for AML2 CD34^+^/CD38-blasts (Figure 5A).

**Figure 3.**
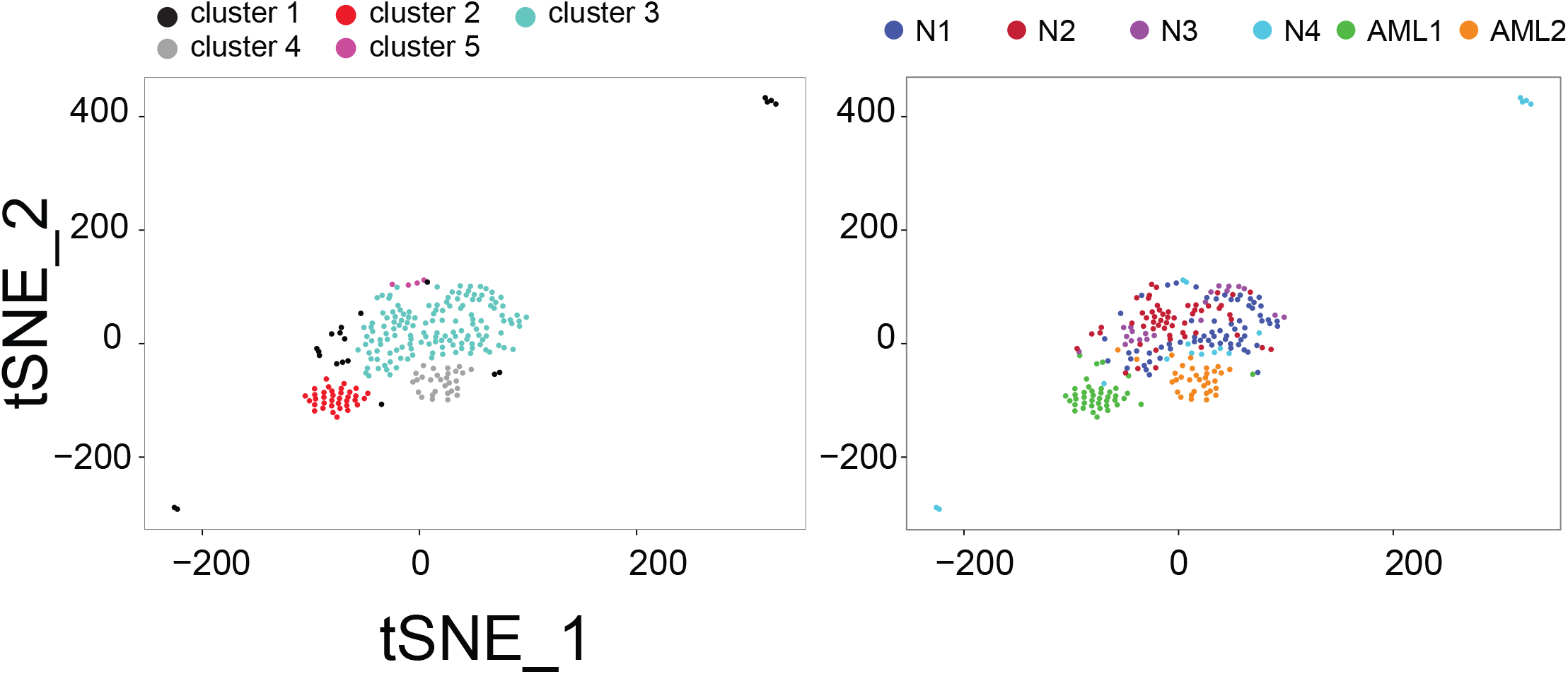
CD33^+^/CD34^+^/CD38^−^ single-cell transcription profiles distinguish leukemic and normal cells. t-SNE map with cells colored by cluster identities (left plot) or by individuals (right plot). Cells were classified into 5 clusters according to their expression patterns using the SEURAT algorithm with the default parameters. Cells that could not be assigned were placed by default into cluster 1.

**Figure 4.**
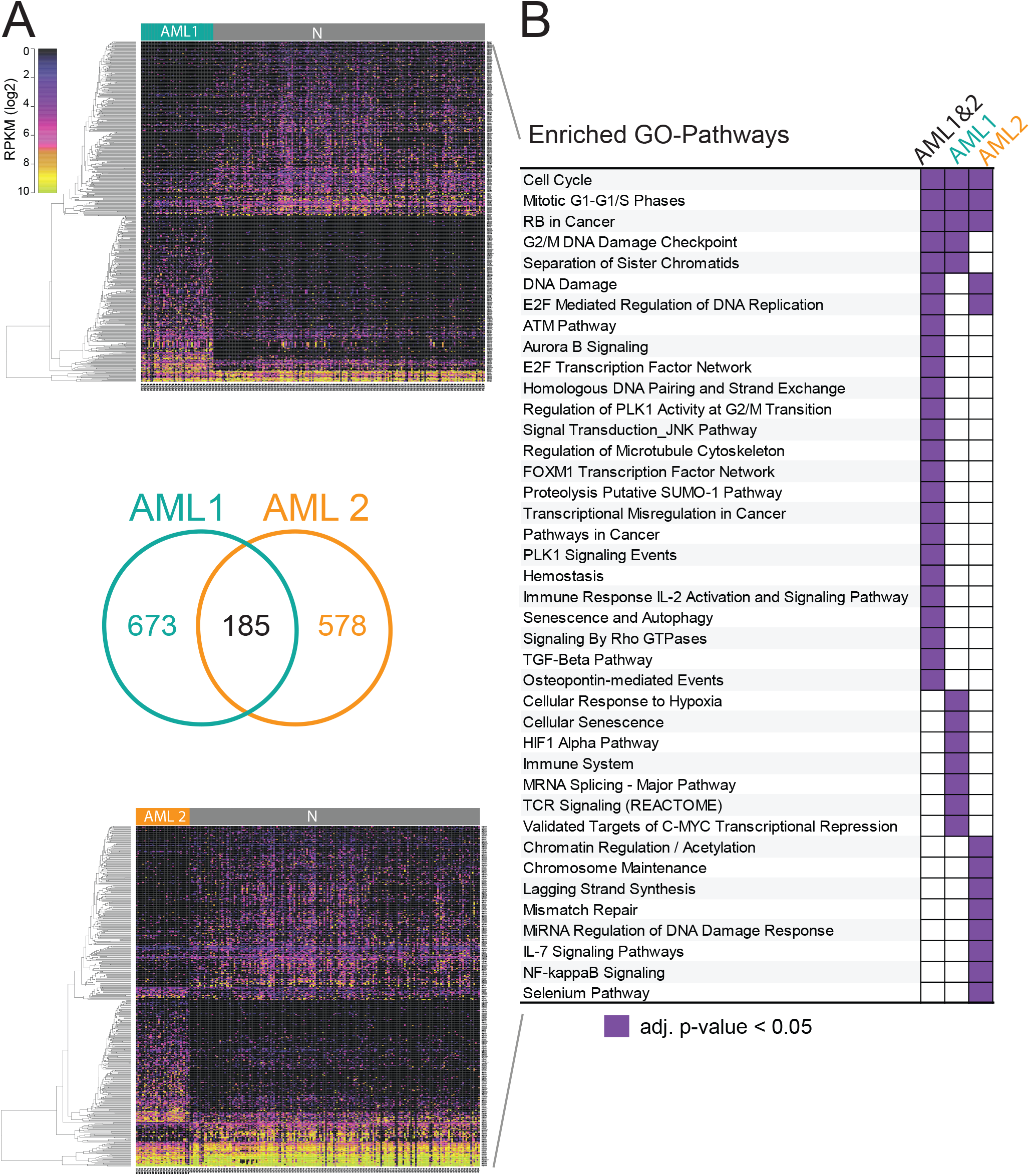
Differential gene expression in CD34^+^/CD38^−^ blasts reveals enriched “stemness”-related signaling pathways compared to non-AML cells. **(A)** Heat maps of the differentially expressed genes (D^3^E analysis) discriminating N cells from AML cells. Genes are shown in rows. Log2-transformed RPKM values are indicated in the color map (refer to the supplementary data, Table S2 and S3). The Venn diagram indicates the number of genes differentially expressed between N and AML1 samples (858 genes) and between N and AML2 samples (763 genes). **(B)** GO term analysis of differentially expressed genes between AML and N samples (GeneAnalytics tool, adj. p-value < 0.05). The purple color indicates the significant enrichment of gene ontology pathways in AML1 (673 genes), AML2 (578 genes) and AML1 & 2 (185 genes).

**Figure 5.**
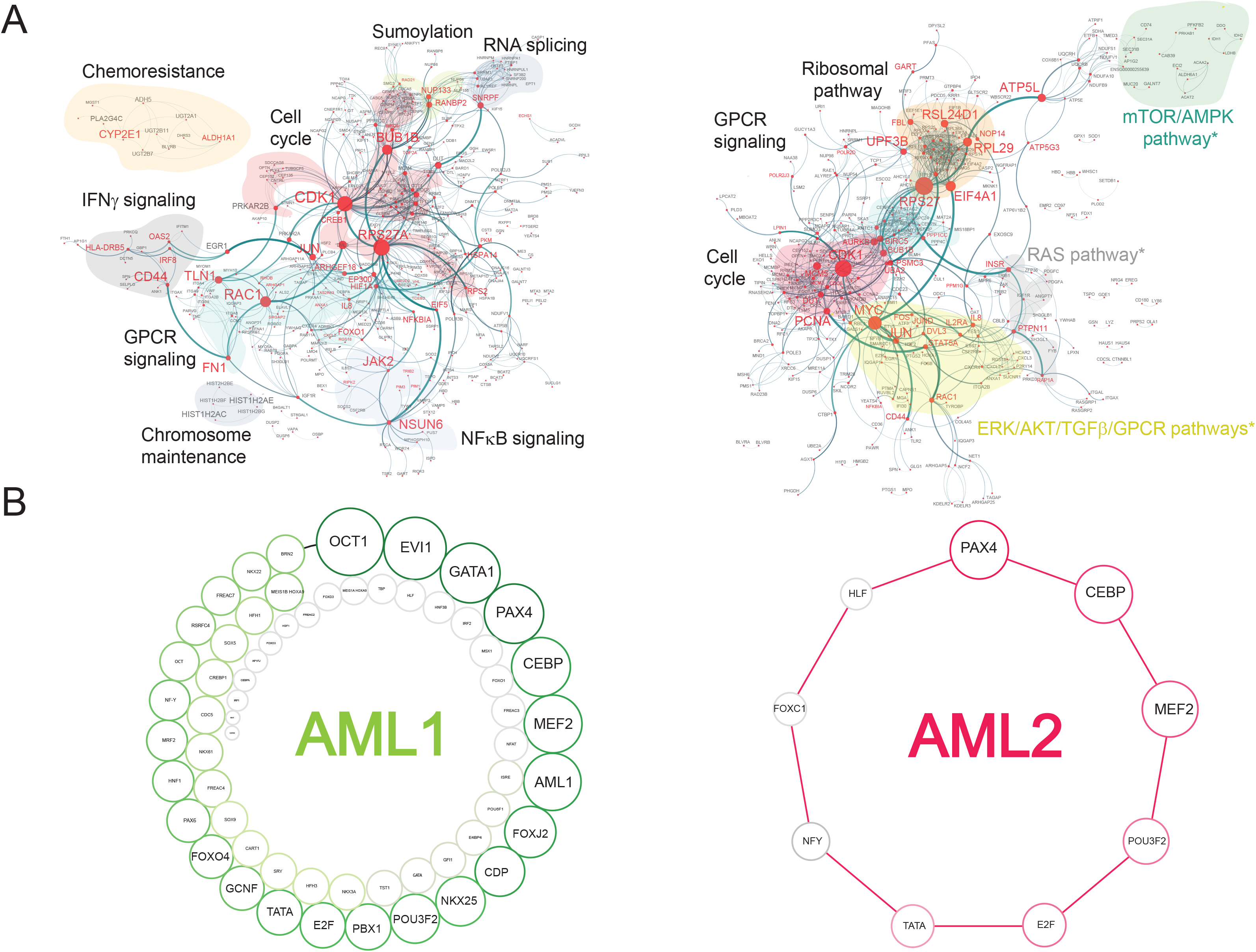
A core set of TFs define a gene-based network associated with disease and stemness-related signaling pathways. **(A)** Interaction map of differentially expressed genes (D^3^E analysis) for AML1 (left) and AML2 (right). Relevant signaling pathways are highlighted. Genes representing the main nodes are colored in red. * indicates pathways known to be perturbed by FL3-ITD mutation. **(B)** Graphic representation of TFBs enriched in differentially expressed gene sets for AML1 (left) and AML2 (right). Circles are sized in proportion to the number of differentially expressed genes enriched for the labelled TFBs.

Overall, our single-cell transcriptome analysis conducted on CD33^+^/CD34^+^/CD38^−^ cells from two AML samples provided valuable information on the nature of the AML. The effects of somatic mutations on the cancer cells are detectable providing valuable insight into disease-associated and patient-specific gene networks.

### Core set of transcription factors are co-activated in leukemic CD34^+^/CD38^−^ cells

To further investigate the regulatory networks of leukemic CD34^+^/CD38^−^ cells, we investigated whether previously identified differentially expressed genes in the AML1 and AML2 cells were co-regulated by a set of transcription factors. Therefore, we evaluated the enrichment of transcription factor binding sites (TFBs) associated with those differentially expressed gene sets (refer to the Materials and Methods). Of the 763 significantly differentially expressed genes in the CD34^+^/CD38^−^ AML2 cells, nine TFBs (PAX4, CEBP, MEF2, POU3F2, E2F, TATA, NFY, FREAC3 (FOXC1) and HLF) were significantly enriched (adjusted p-value < 0.05), and of the 858 differentially expressed genes in the CD34^+^/CD38^−^ AML1 cells, 62 TFBs (the nine TFBs associated with the AML2 dataset as well as other relevant TBFs, such as OCT1, GATA1, EVI1 and MEF2 (Figure 5B)) were significantly enriched (adjusted p-value < 0.05). Interestingly, nine common TFBs are found in common between the CD34^+^/CD38^−^ blasts of both AML samples. This result suggests a relationship among leukemic CD33^+^/CD34^+^/CD38^−^ cells such as a joint transcription program or a shared cellular identity.

### Transcription profiles of leukemic CD34^+^/CD38^−^ single-cells can identify genes associated with putative survival outcomes in the AML-TCGA cohort

Reports from AML patients have indicated that the gene expression signatures of leukemic blasts at diagnosis have prognostic significance [11, 43]. Thus, we hypothesized that these signatures could be detected in leukemic CD33^+^/CD34^+^/CD38^−^ single-cell transcription profiles. RNA-seq data and clinical outcome data of 163 individuals with AML accessible on The Cancer Genome Atlas (TCGA) public website [44] were examined, and then the associations among 1675 differentially expressed genes from our study were assessed against the clinical outcomes in the AML-TCGA cohort. Specifically, we ranked the genes according to the p-values derived from a univariate Cox regression. Thirty-six genes from the AML2 gene set and 22 genes from the AML1 gene set were identified as significant for overall survival based on a comparison of the patients in the top 50% of gene expression against those in the bottom 50% of gene expression (p-value ≤ 0.05) (refer to the supplementary data, Table S5). Five genes (*MPO, ITGAX, RUFY3, FEM1C*, and *HSF2*) were common between the AML1 and AML2 gene sets. The effect of the relative expression of selected genes on the survival of AML-TCGA patients is shown in Kaplan-Meier plots in Figure 6. Interestingly, the overall expression of myeloperoxidase (*MPO*) was higher in the normal CD33^+^/CD34^+^/CD38^−^ single cells compared with the AML cells (FC N/AML2=24.77, p-value=3.92x10^−10^; FC N/AML1=32, p-value=3.97x10^−9^), whereas high levels of *MPO* transcripts were significantly associated with improved survival in the AML-TCGA samples (p-value=0.0042) (Figure 6). *MPO* is a myeloid lineage marker that has been described as a potential prognostic factor for AML [45–48]. This analysis identified genes whose single-cell expression in leukemic cells was associated with the survival of AML-TCGA patients (refer to the supplementary data, Table S5). Moreover, we found an enrichment of previously published prognostic gene signatures [12, 49] in single leukemic CD34^+^/CD38^−^ cells (refer to the supplementary data, Figure S5). Collectively, these findings indicate that the transcriptional landscape of leukemic CD33^+^/CD34^+^/CD38^−^ cells contains putative relevant disease outcome information and the single-cell transcriptomic strategy presented here is of potential use in further prognostic evaluations.

**Figure 6.**
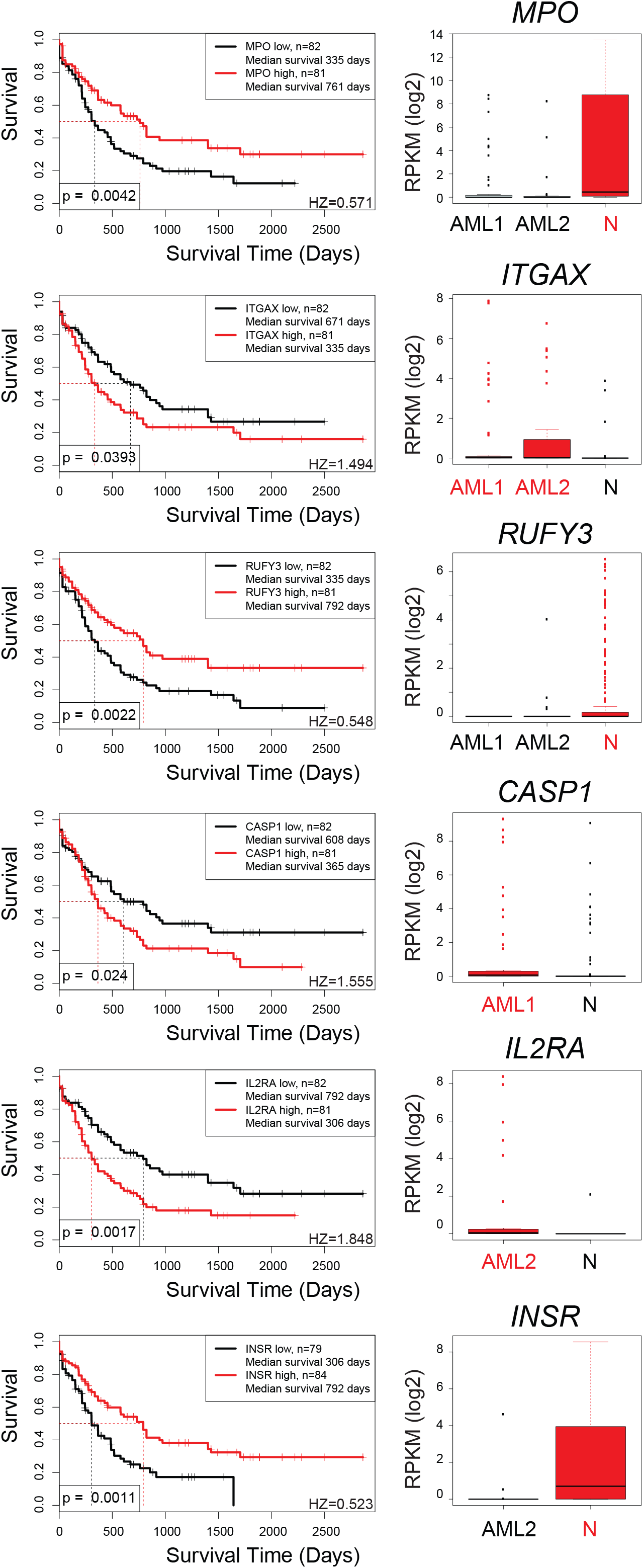
CD33^+^/CD34^+^/CD38^−^ single-cell transcription profiles identify genes significantly associated with survival outcomes in the AML-TCGA cohort (adj. p-value < 0.05). (Left) Kaplan-Meier survival curves for AML-TCGA patients stratified based on transcript levels. Patients were divided into two groups: 50% lowest gene expression and 50% highest gene expression. P-values were calculated using a univariate Cox regression analysis. (Right) Box-plots displaying the distribution of transcript levels (log2-transformed RPKM) in AML1, AML2 and N single-cell sample sets. The groups of cells with the highest significant expression are shown in red (p-value < 0.05) (refer to the supplementary data, Table S5). HZ: hazard ratio.

## DISCUSSION

Utilizing a single-cell RNA sequencing approach, we examined for the first time the transcriptional profile of normal and malignant individual CD33^+^/CD34^+^/CD38^+/−^.cells. We focused our analysis on leukemic CD33^+^/CD34^+^/CD38^−^ blasts from two patients with AML by comparing their transcriptional profile to more differentiated CD33^+^/CD34^+^/CD38^+^blasts and normal CD33^+^/CD34^+^/CD38^−^ cells.

Our findings revealed transcriptional regulatory networks associated with cell cycle features of normal and tumor CD33^+^/CD34^+^/CD38^−^ cells. The CD33^+^/CD34^+^/CD38’ AML cells displayed a notable deregulation of the cell cycle and cellular differentiation programs compared with the CD34^+^/CD38^−^ non-AML cells. Indeed, different cyclin-dependent kinases and cell cycle regulatory proteins were activated in the CD34^+^/CD38^−^ blasts and myeloid differentiation processes were repressed, which also occurs in leukemogenic cells [50]. Additionally, our analysis revealed deregulation of the mTORC/AMPK, ERK/MAPK, AKT, STAT3/5, IFNγ, GPRC and NF-κB pathways in single leukemic CD34^+^/CD38^−^ cells. Altogether, our results suggest that leukemic CD34^+^/CD38^−^ cells undergo abnormal myeloid differentiation to a controlled proliferative state and exhibit the ability to regulate their self-renewal potential.

Although gene ontology analyses do not provide a clear description of the mechanisms that regulate the proliferation and differentiation of leukemic cells, they can be used to validate observations of interactive networks specific to each AML. Alternatively, the study of groups of deregulated genes and associated pathways enables the identification of transcription factors that are co-activated in leukemic cells. This approach can lead to the discovery of new features associated with a stem-like phenotype, such as self-renewal, survival, myeloid differentiation and cancer progression mechanisms. Our findings showed that the highly connected core network circuit in CD34^+^/CD38^−^ blasts (AML1 and AML2) is controlled by a set of transcription factors: C/EBP (CCAAT enhancer binding protein alpha), a main determinant of myeloid differentiation [51–53]; POU family members Pit-1, Oct1/2, Unc-86 [54] [55] and POU3F2 [56], which are known to drive self-renewal and stress resistance; FOXC1, which controls stem cell function [57]; MEF2, which is known for its role in hematopoietic cell differentiation [58, 59]; NF-Y, which is a regulator of HSC proliferation and survival [60] and HLF, which is known for its self-renewal capacity *in vitro* in HSCs and progenitors [61]. Additionally, GATA1, EVI1 and AML1 (RUNX1) were exclusively activated in CD34^+^/CD38^−^ cells from patient AML1. The role of these transcription factors in stem cell self-renewal and leukemogenesis has also been well established [62–66].

We also reported the activation of several pathways related to oncogenic events specific to each type of AML. Remarkably, cells from patient AML2 harbored a FLT3-ITD oncogene mutation and exhibited the specific activation of the PI3K/AKT/mTOR, RAS/MAPK and Janus kinase 2 (JAK2)/STAT5 pathways as a consequence of the constitutive activation of the FLT3 receptor [67]. As a result, the clustering analysis performed on the single-cell transcriptomes was sufficient to discriminate normal cells from leukemic cells as well as AML1 cells from AML2 cells. This study is consistent with the hypothesis that CD34^+^/CD38 populations transcriptionally differ depending upon specific mutations that generate heterogeneous subsets of blasts with specific deregulated pathways. Importantly, transcriptional analyses at the single-cell level, such as the analysis performed in this study, can monitor the effects of oncogenic events on signaling pathways and provide information on the minimal residual disease and potential outcome of therapies.

In addition, differentially deregulated genes in the CD34^+^/CD38^−^ AML cells were found to be significantly associated with patient survival compared with the CD34^+^/CD38^−^ non-AML cells. Five genes (*MPO, ITGAX, RUFY3, FEM1C*, and *HSF2*) emerged from our study as putative candidates implicated in patient survival. Previous works have associated the percentage of MPO-positive cells with positive prognosis in patients with CD34^+^/CD38^−^ AML [45–48], thus supporting our findings. *ITGAX* (or *CD11c*) is a myeloid marker regulated by the non-coding RNA HOX antisense intergenic RNA myeloid 1 (*HOTAIRM1*), which is in the HOXA cluster [68]. Our results are consistent with the reported association between *HOTAIRM1* expression and low patient survival in intermediate-risk AML [69]. A promising marker is the insulin receptor gene (*INSR* or IR), a growth factor receptor tyrosine kinase that is downregulated in CD33^+^/CD34^+^/CD38^−^ AML2 cells compared with non-AML cells. This gene was significantly associated with poor prognosis when expressed at lower levels in patients with AML in our study. *INSR* is a key regulator of the PI3K/AKT/mTOR signaling pathway [70] and known to regulate carbohydrate metabolism [71]. Our results as well as the results of other groups [72] indicate that this central pathway is strongly deregulated in leukemic CD34^+^/CD38^−^ cells. The relationship found between *INSR* transcript levels and AML patient survival is consistent with the hypothesis that the PI3K/AKT/mTOR signaling pathway plays a crucial role in leukemia-initiating cells and might represent a putative novel gene expression biomarker of survival. Overall, transcriptional regulatory circuits can be revealed in leukemic CD33^+^/CD34^+^/CD38^−^ cells using single-cell RNA-seq. This approach can be used to study cellular transcriptional states of rare cancer cells at an unprecedented resolution. Notably, single-cell analyses provide a tool to study relationship between differentially expressed genes in leukemia and patient prognosis as described in the AML-TCGA database.

## CONCLUSIONS

This study demonstrated the power and the unique contribution of single-cell RNAseq to explore rare cancer cell types by developing gene expression signatures and revealing the disease status of each patient. Future studies that integrate our current workflow on a larger cohort of AML individuals and cells would enable the identification of new significant gene biomarkers for AML natural history and treatment monitoring and choices.

## MATERIALS AND METHODS

### Samples

Fresh bone marrow samples were obtained from patients of the Geneva Hematology Service in accordance with institutional guidelines and after approval by the ethics committee of the Geneva University Hospital. Written informed consent was obtained from all patients. Two patients were diagnosed with AML: Patient AML1 (83-year-old male), who had AML M0 with a normal karyotype without FLT3-ITD and NPM1 mutations; and Patient AML2 (52-year-old male), who had AML M1 with a normal karyotype harboring FLT3-ITD and NPM1 mutations. CEBPA mutations were absent and WT1 gene was found overexpressed in both patients. The other four patients underwent a bone marrow aspiration in a diagnostic work-up and were found to exhibit normal bone marrow after morphologic, flow cytometric, and karyotypic analyses.

### Cell staining and flow cytometry

Cell suspensions were stained with the following antibodies: CD45=FITC (Dako), CD34=PE (Beckman Coulter), CD38=BV421 (BD Biosciences), CD19=APC (Beckman Coulter), and CD33=PE Cy7 (Beckman Coulter). Single cells from the CD34^+^, CD38^−^, CD33^−/+^, CD19^−^ normal myeloid HSC or LSC sub-populations were sorted using a cell sorter (MoFlo Astrios cell sorter, Beckman Coulter).

### Single-cell RNA-sequencing

Single-cell captures were performed for the six patient samples on the C1 Single-Cell Auto Prep System (Fluidigm) using the cell load script 1772x/1773x. Approximately 2500–5000 CD33^+^/CD34^+^/CD38^+^ live cells were directly sorted into the assay well of a primed microfluidic array (C1 Single-Cell Auto Prep Array for mRNA-seq, 5–10 μm, 96 chambers, Fluidigm) following the manufacturer’s protocol. For further analysis, only chambers containing a single cell by visual inspection were considered, and chambers containing debris, multiple or damaged cells and empty chambers were discarded [73]. cDNA preparations were performed on a C1 Single-Cell Array for mRNA-seq using the C1 Single-Cell Auto Prep System (Fluidigm). A SMARTer Ultra Low RNA Kit for Illumina Sequencing (version 2, Takara Clontech) was used for cell lysis and cDNA synthesis following the manufacturer’s protocol [73]. One capture array per patient was analyzed for a total of six single-cell experiments.

Because only one C1 single-cell array can be processed at a time, we modified our single-cell capture protocol to enable parallel analysis of the CD38^+^ and CD38^−^ cells from the AML2 sample. The CD33^+^/CD34^+^/CD38^−^- and CD33^+^/CD34^+^/CD38^+^-stained cells were sorted directly into two 96-well PCR plates (1 cell per well) containing 2 μl of 1x Dulbecco’s phosphate-buffered saline (Gibco, Life Technologies). Then, cDNA synthesis was performed using a SMART-Seq v4 Ultra Low Input RNA Kit for Sequencing (Takara Clontech) following the manufacturer’s protocol.

For all of the samples, the cDNA quality was assessed on a 2100 Bioanalyzer (Agilent) using high sensitivity DNA chips (Agilent). Subsequently, the cDNA was quantified using a Qubit dsDNA BR Assay Kit (Invitrogen). Sequencing libraries were prepared with 0.3 ng of pre-amplified cDNA using a Nextera XT DNA Kit (Illumina) according to the manufacturer’s instructions [73]. Libraries were sequenced on an Illumina HiSeq 2000 machine as single-end 100 bp reads with a multiplexing of 12 cells per lane of the flow cell. A total of 359 single cells were sequenced: 49 single-cells for AML1, 37 for AML2, 77 for N1, 68 for N2, 34 for N3, and 46 for N4. For AML2, an additional 24 CD33^+^/CD34^+^/CD38^−^ single cells and 24 CD33^+^/CD34^+^/CD38^+^ single cells were sequenced.

### Sequencing of bulk RNA

Libraries were prepared from bulk RNA of 300 sorted CD33^+^/CD34^+^/CD38^−^ cells using a SMARTer Ultra Low RNA Kit for Illumina Sequencing (version 2, Takara Clontech) according to the manufacturer’s instructions. Libraries were sequenced on an Illumina HiSeq 2000 machine as 100 bp single-end reads.

### Processing of sequencing data

Alignment and mapping of the sequencing reads were performed against the human genome (GRCh38/hg38) using Bowtie2 (version 2.2.4) [74] and TopHat (version 2.0.11) [75] with the default settings. Cells with fewer than 20% of uniquely mapped reads were filtered out. After quality filtering, 313 single-cell RNA-seq samples were analyzed in this study: 49 AML1, 34 AML2, 74 N1, 66 N2, 27 N3, 17 N4, 23 CD33^+^/CD34^+^/CD38’ AML2, and 23 CD33^+^/CD34^+^/CD38^+^ AML2. The expression levels of transcripts were quantified and normalized as reads per kilobase per million (RPKM) with Gencode annotation (version 21). The following GENCODE categories were considered for the study: TR genes, IG genes, lincRNAs, snoRNAs, miRNAs and protein coding genes. ChrY genes were excluded to avoid gender bias. This process resulted in 32 511 GENCODE genes in total.

### Principal component clustering analysis

A principal component analysis (PCA) was performed using R (version 3.1.3) [76]. A clustering analysis and t-distributed stochastic neighbor embedding (t-SNE) visualization were performed with the Seurat R package (version 1.2) [77]. Default-parameters were used with at least 3 cells per gene and cells with more than 1000 detected genes (RPKM > 1).

### Differential transcript expression analysis

Differentially expressed genes were identified using Discrete Distributional Differential Expression (D(3)E) for single-cells [78] in Python using the Kolmogorov-Smirnov statistical test (p-value < 0.05). In addition, the most relevant genes with a burst frequency calculated by D(3)E (Rf) at an absolute value greater than 2 were selected [79]. Lists of differentially expressed genes are provided in the supplementary data (Tables S1-3).

### Gene ontology and gene set enrichment analysis

DAVID 6.7 [80] and GeneAnalytics [81] were used for the enrichment analysis of pathways. Gene set enrichment analyses were conducted using GSEA software (version 2.2.0) [82], and corrected p-values were assessed using 10 000 permutations of the gene sets. Annotated gene sets from the Molecular Signature Database (MSigDB), including the KEGG database (Kyoto Encyclopedia of Genes and Genomes, http://www.genome.jp/kegg/) and published gene datasets, were used.

### Complex-network analysis

Network modes and edges were generated using the Search Tool for the Retrieval of Interacting Genes (STRING, version 10) [83]. Visual mapping was performed with Cytoscape 3.3.0 [84].

### Transcription factor analysis

An enrichment analysis for TFBs was performed using the UCSC_TFBS track option in the functional annotation tool of DAVID 6.7 [85, 86]. The UCSC_TFBS in DAVID is based on the tfbsConsSites table of the UCSC genome browser.

### Survival analysis

Gene expression and clinical information were derived from The Cancer Genome Atlas (TCGA) acute myeloid leukemia dataset (n=173) [87]. Significant differences in survival rates were evaluated using a univariate Cox regression models (p-value < 0.05). The hazard ratio was reported in comparison to the gene low-expression group. The survival curves were visualized using Kaplan-Meier plots.

## AUTHOR CONTRIBUTIONS

Conceptualization: C.B, T.M. and S.E.A; Methodology: C.B, T.M., A.S., E.F., P.R., P.C. and M.P.L.; Investigation: C.B., T.M., A.S. and S.E.A.; Writing of the Original Draft: C.B., A.S. and T.M.; Writing, Review and Editing: C.B., T.M., S.E.A., A.S., E.F., P.R., P.C. and M.P.L.; Funding Acquisition: C.B. and T.M.; Patient Recruitment: T.M.; and Patient Supervision: C.B. and T.M.

## ACCESSION CODES

RNA sequencing data were deposited in the European Genome-phenome Archive (EGA, https://www.ebi.ac.uk/ega/) for controlled access; the study accession number is (*to be determined*).

## ACKNOWLEDGMENTS

The authors thank the Flow Cytometry Facility and the iGE3 genomics platform of the University of Geneva. The authors also thank Sylvie Ruault-Jungblut for expert technical assistance. This study was supported by a grant from the Dr. Henri Dubois-Ferrière Dinu Lipatti Foundation (http://www.dfdl.org). The funders had no role in the data collection, study design and analysis, decision to publish, or preparation of the manuscript. Calculations were performed at the Vital-IT Center (http://www.vital-it.ch) for high-performance computing of the SIB Swiss Institute of Bioinformatics.

The authors declare no financial conflict of interests.

## SUPPLEMENTAL INFORMATION

**Figure S1.**
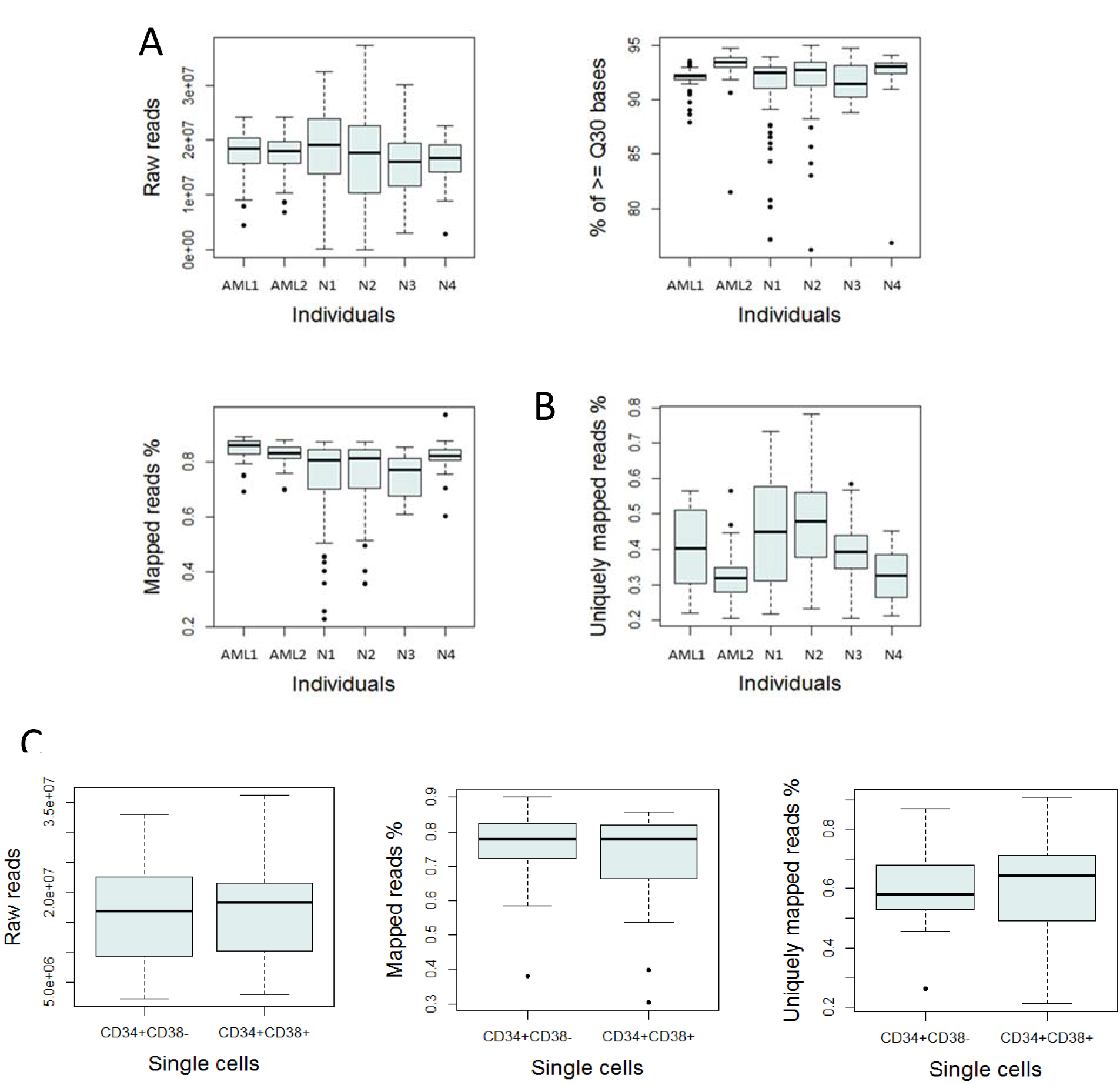
Statistics of reads mapping. Related to figure 1. **(A)** Boxplots summarizing the number of raw reads, percentage of reads with a quality over Q30, the percentage of mapped reads over the different single-cell samples (311 cells in total). **(B)** Boxplot representing the percentage of uniquely mapped reads after filtering out cells with less than 20% of mapped reads (267 cells in total). **(C)** Boxplots summarizing the number of raw reads (left), the percentage of mapped reads (middle), the percentage of uniquely mapped reads after filtering out cells with less than 20% of mapped reads (right) over AML2 CD33+CD34+CD38+ and AML2 CD33+CD34+CD38-single-cell samples (46 cells in total).

**Figure S2.**
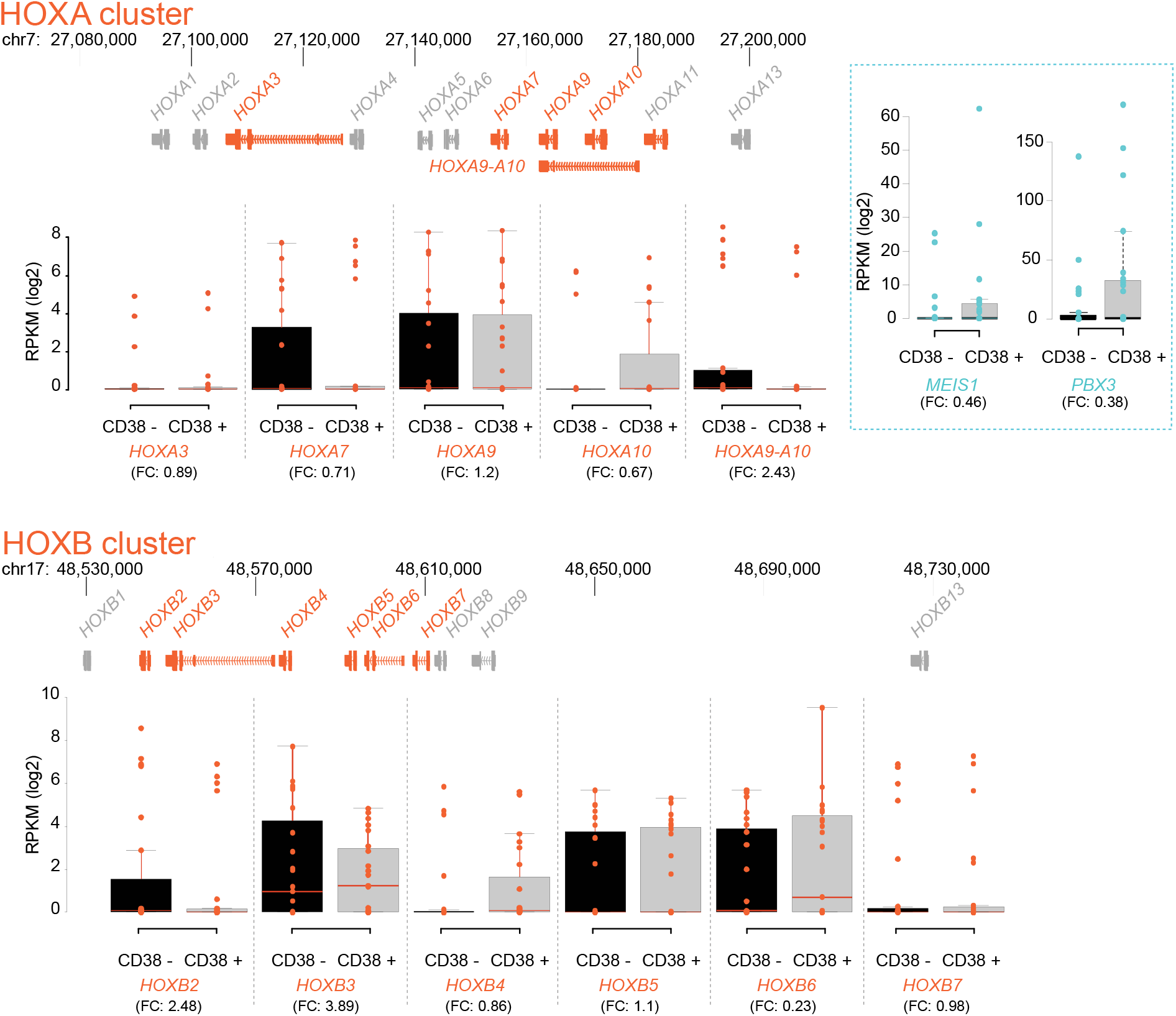
Genomic organization of the *HOX A* and *HOX B* cluster on chromosome 7 and 17 respectively. Related to figure 2 and table S1. Maps are based on UCSC genome browser (hg38). Expressed genes are colored in orange. Box plots of Log2 transformed RPKM of detected *HOX* genes. *MEIS 1* and *PBX3*, co-factors of *HOX* genes, are plotted in a separate dashed turquoise panel on the right. FC: average fold change of CD38−/CD38+.

**Figure S3.**
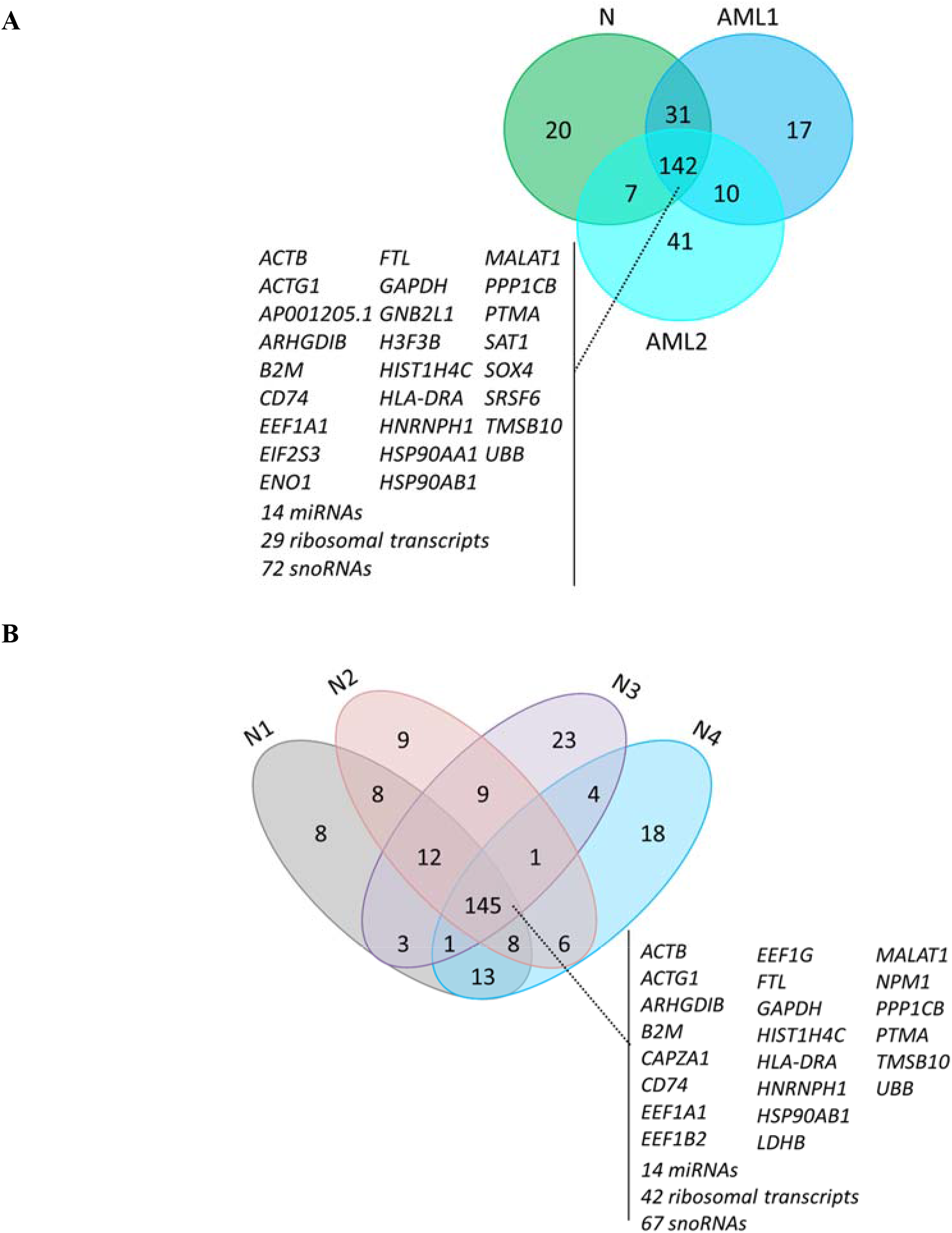
The top 200 most expressed genes. Related to figure 3. Venn diagram of the top 200 most expressed expressed **(A)** in AML and non-AML single-cell samples and **(B)** in 4 non-AML single-cell samples.

**Figure S4.**
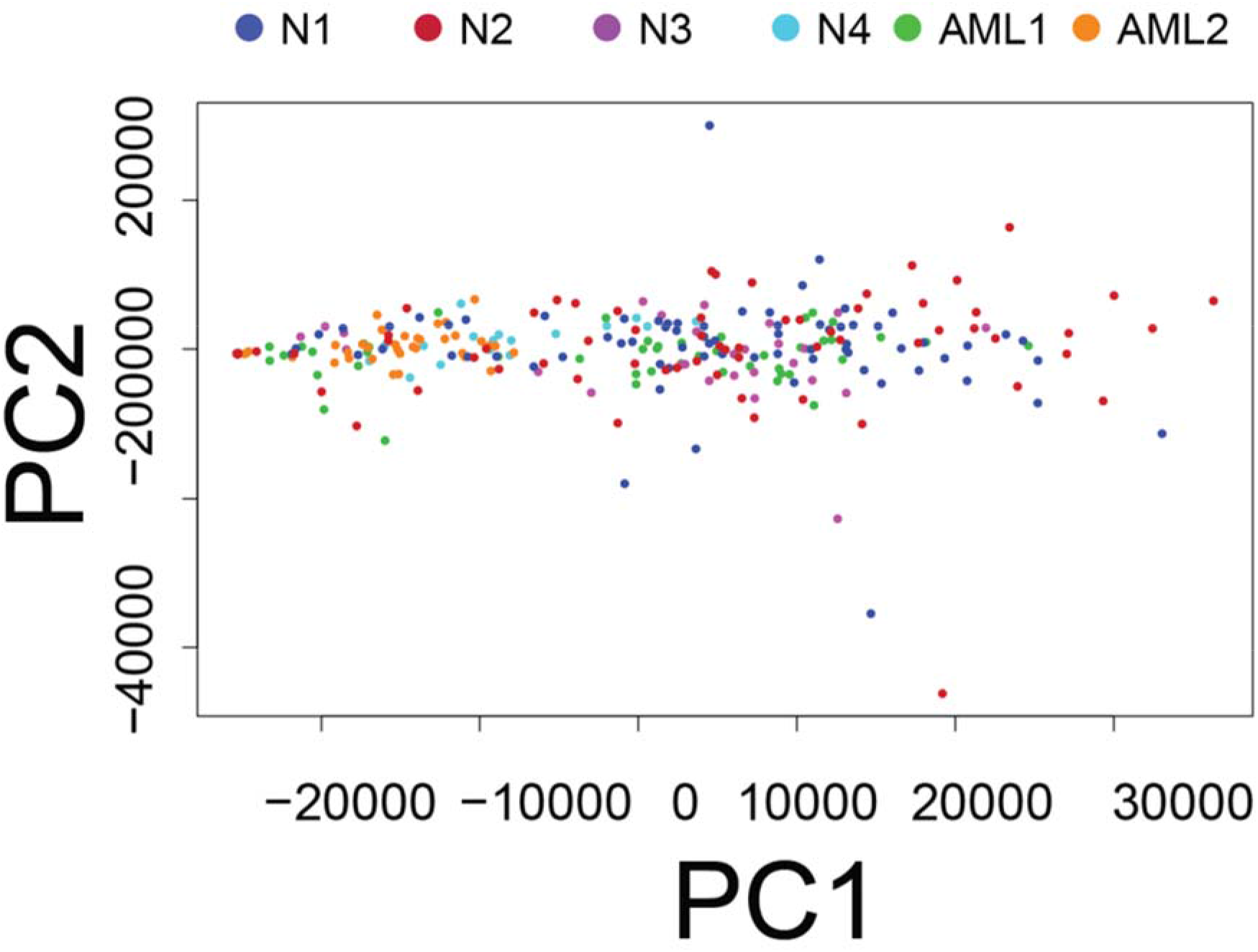
PCA analysis. Related to figure 3. Plot showing the first two principal components of 267 normal and leukemic CD33+/CD34+/CD38− cells. We observed that the 267 transcriptome profiles are relatively dispersed. The first two principal components, respectively, account for 69% and 2,8% of the total variability in gene expression.

**Figure S5.**
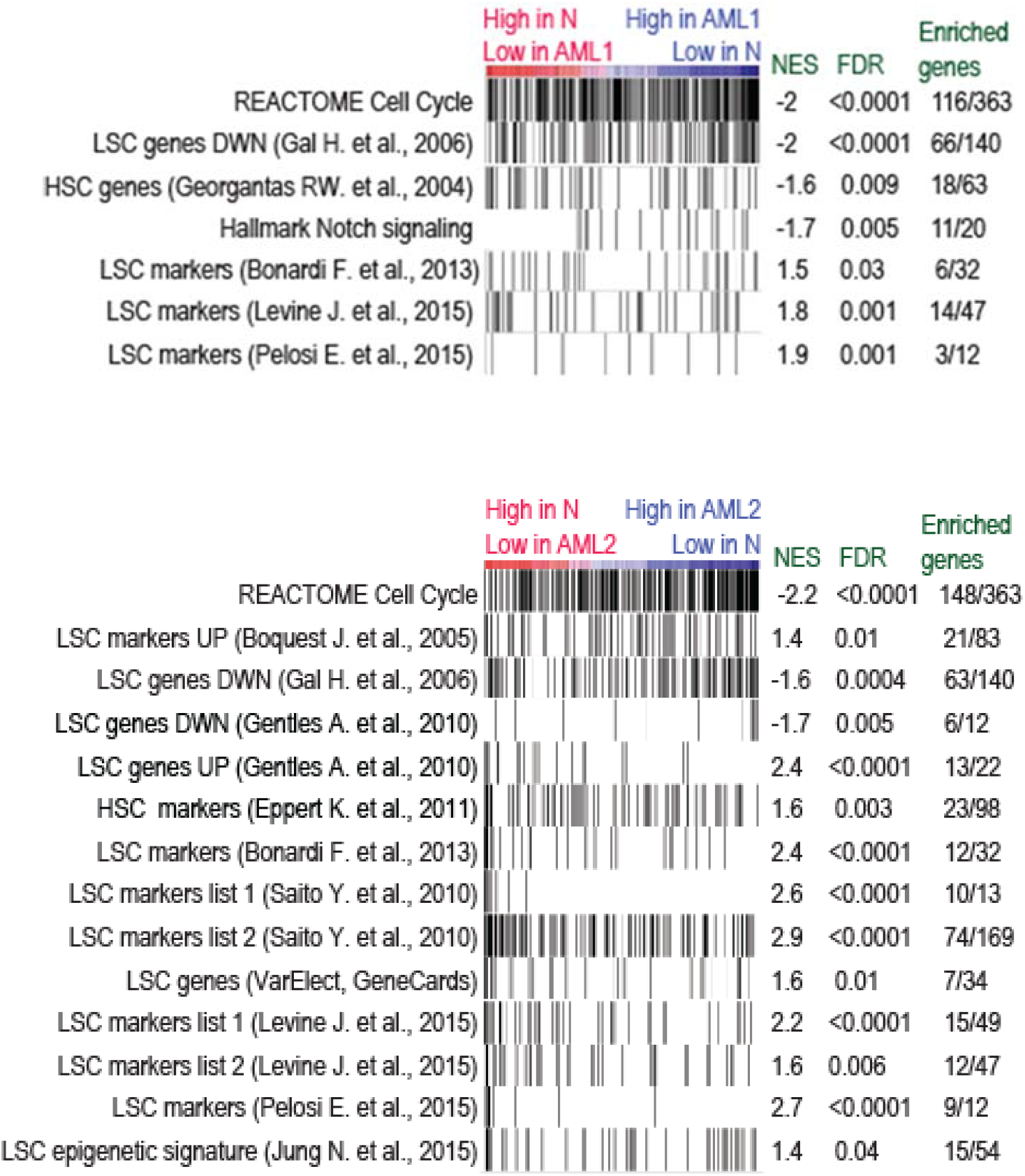
Signature enrichment plots from GSEA analysis using different genes target lists. Related to figure 4 and table S4. Black vertical bars are ordered genes according to their fold change expression. Values indicated normalized enrichment score (NES), FDR adjusted p-value and number of significant enriched genes out of the total tested genes. Tested gene lists are detailed in Table S4. Results for non-AML versus AML1 single-cell samples are showed in top panel and non-AML versus AML2 single-cell samples in the bottom panel.

**Figure S6.**
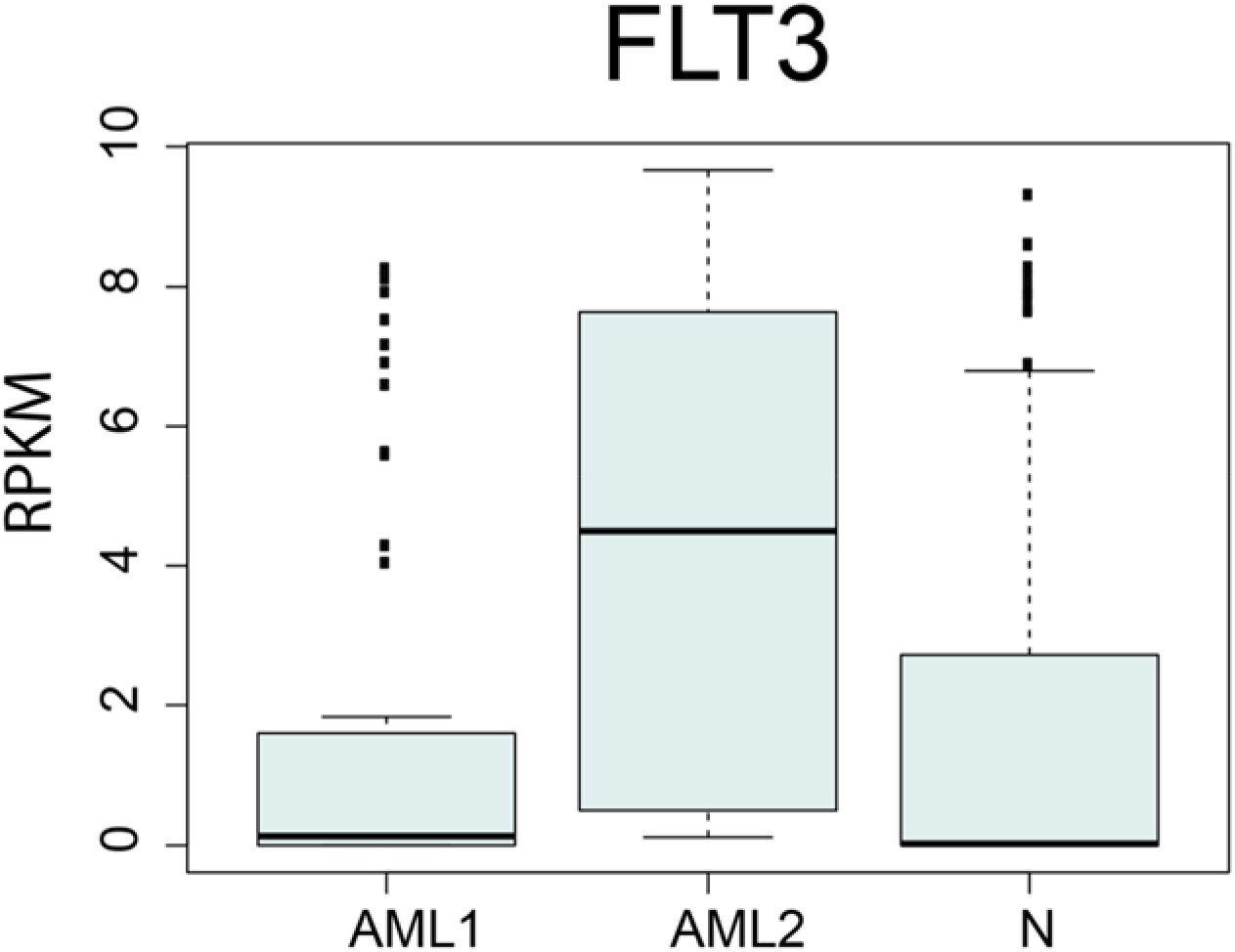
Gene expression of FLT3 transcript. Related to figure 5. Boxplot of the Log(2) transformed RPKM data of *FLT3* gene.

## REFERENCES

1. Lowenberg B, Downing JR, Burnett A: Acute myeloid leukemia. N Engl J Med 1999, 341:1051–1062.

2. Dohner H, Weisdorf DJ, Bloomfield CD: Acute Myeloid Leukemia. N Engl J Med 2015, 373:1136–1152.

3. Bonnet D, Dick JE: Human acute myeloid leukemia is organized as a hierarchy that originates from a primitive hematopoietic cell. Nat Med 1997, 3:730–737.

4. Lapidot T, Sirard C, Vormoor J, Murdoch B, Hoang T, Caceres-Cortes J, Minden M, Paterson B, Caligiuri MA, Dick JE: A cell initiating human acute myeloid leukaemia after transplantation into SCID mice. Nature 1994, 367:645–648.

5. Dick JE: Stem cell concepts renew cancer research. Blood 2008, 112:4793–4807.

6. Schepers K, Campbell TB, Passegue E: Normal and leukemic stem cell niches: insights and therapeutic opportunities. Cell Stem Cell 2015, 16:254–267.

7. Dick JE: Acute myeloid leukemia stem cells. Ann N Y Acad Sci 2005, 1044:1–5.

8. Sarry JE, Murphy K, Perry R, Sanchez PV, Secreto A, Keefer C, Swider CR, Strzelecki AC, Cavelier C, Recher C, et al: Human acute myelogenous leukemia stem cells are rare and heterogeneous when assayed in NOD/SCID/IL2Rgammac-deficient mice. J Clin Invest 2011, 121:384–395.

9. Forsberg EC, Passegue E, Prohaska SS, Wagers AJ, Koeva M, Stuart JM, Weissman IL: Molecular signatures of quiescent, mobilized and leukemia-initiating hematopoietic stem cells. PLoS One 2010, 5:e8785.

10. Bonardi F, Fusetti F, Deelen P, van Gosliga D, Vellenga E, Schuringa JJ: A proteomics and transcriptomics approach to identify leukemic stem cell (LSC) markers. Mol Cell Proteomics 2013, 12:626–637.

11. Gentles AJ, Plevritis SK, Majeti R, Alizadeh AA: Association of a leukemic stem cell gene expression signature with clinical outcomes in acute myeloid leukemia. JAMA 2010, 304:2706–2715.

12. Eppert K, Takenaka K, Lechman ER, Waldron L, Nilsson B, van Galen P, Metzeler KH, Poeppl A, Ling V, Beyene J, et al: Stem cell gene expression programs influence clinical outcome in human leukemia. Nat Med 2011, 17:1086–1093.

13. van Rhenen A, Feller N, Kelder A, Westra AH, Rombouts E, Zweegman S, van der Pol MA, Waisfisz Q, Ossenkoppele GJ, Schuurhuis GJ: High stem cell frequency in acute myeloid leukemia at diagnosis predicts high minimal residual disease and poor survival. Clin Cancer Res 2005, 11:6520–6527.

14. Mikkola HK, Radu CG, Witte ON: Targeting leukemia stem cells. Nat Biotechnol 2010, 28:237–238.

15. Kolodziejczyk AA, Kim JK, Svensson V, Marioni JC, Teichmann SA: The technology and biology of single-cell RNA sequencing. Mol Cell 2015, 58:610–620.

16. Shapiro E, Biezuner T, Linnarsson S: Single-cell sequencing-based technologies will revolutionize whole-organism science. Nat Rev Genet 2013, 14:618–630.

17. Grun D, Lyubimova A, Kester L, Wiebrands K, Basak O, Sasaki N, Clevers H, van Oudenaarden A: Single-cell messenger RNA sequencing reveals rare intestinal cell types. Nature 2015, 525:251–255.

18. Trapnell C: Defining cell types and states with single-cell genomics. Genome Res 2015, 25:1491–1498.

19. Shalek AK, Satija R, Adiconis X, Gertner RS, Gaublomme JT, Raychowdhury R, Schwartz S, Yosef N, Malboeuf C, Lu D, et al: Single-cell transcriptomics reveals bimodality in expression and splicing in immune cells. Nature 2013, 498:236–240.

20. Marinov GK, Williams BA, McCue K, Schroth GP, Gertz J, Myers RM, Wold BJ: From single-cell to cell-pool transcriptomes: stochasticity in gene expression and RNA splicing. Genome Res 2014, 24:496–510.

21. Buettner F, Natarajan KN, Casale FP, Proserpio V, Scialdone A, Theis FJ, Teichmann SA, Marioni JC, Stegle O: Computational analysis of cell-to-cell heterogeneity in single-cell RNA-sequencing data reveals hidden subpopulations of cells. Nat Biotechnol 2015, 33:155–160.

22. Stegle O, Teichmann SA, Marioni JC: Computational and analytical challenges in single-cell transcriptomics. Nat Rev Genet 2015, 16:133–145.

23. Waugh DJ, Wilson C: The interleukin-8 pathway in cancer. Clin Cancer Res 2008, 14:6735–6741.

24. Schinke C, Giricz O, Li W, Shastri A, Gordon S, Barreyro L, Bhagat T, Bhattacharyya S, Ramachandra N, Bartenstein M, et al: IL8-CXCR2 pathway inhibition as a therapeutic strategy against MDS and AML stem cells. Blood 2015, 125:3144–3152.

25. Hyka-Nouspikel N, Lemonidis K, Lu WT, Nouspikel T: Circulating human B lymphocytes are deficient in nucleotide excision repair and accumulate mutations upon proliferation. Blood 2011, 117:6277–6286.

26. Beerman I, Seita J, Inlay MA, Weissman IL, Rossi DJ: Quiescent hematopoietic stem cells accumulate DNA damage during aging that is repaired upon entry into cell cycle. Cell Stem Cell 2014, 15:37–50.

27. Nouspikel T: DNA repair in differentiated cells: some new answers to old questions. Neuroscience 2007, 145:1213–1221.

28. Gal H, Amariglio N, Trakhtenbrot L, Jacob-Hirsh J, Margalit O, Avigdor A, Nagler A, Tavor S, Ein-Dor L, Lapidot T, et al: Gene expression profiles of AML derived stem cells; similarity to hematopoietic stem cells. Leukemia 2006, 20:2147–2154.

29. Georgantas RW, 3rd, Tanadve V, Malehorn M, Heimfeld S, Chen C, Carr L, Martinez-Murillo F, Riggins G, Kowalski J, Civin CI: Microarray and serial analysis of gene expression analyses identify known and novel transcripts overexpressed in hematopoietic stem cells. Cancer Res 2004, 64:4434–4441.

30. Wang Y, Krivtsov AV, Sinha AU, North TE, Goessling W, Feng Z, Zon LI, Armstrong SA: The Wnt/beta-catenin pathway is required for the development of leukemia stem cells in AML. Science 2010, 327:1650–1653.

31. Ramsay RG, Gonda TJ: MYB function in normal and cancer cells. Nat Rev Cancer 2008, 8:523‐534.

32. Baker SJ, Ma’ayan A, Lieu YK, John P, Reddy MV, Chen EY, Duan Q, Snoeck HW, Reddy EP: B-myb is an essential regulator of hematopoietic stem cell and myeloid progenitor cell development. Proc Natl Acad Sci U S A 2014, 111:3122–3127.

33. Jiang L, Li J, Song L: Bmi-1, stem cells and cancer. Acta Biochim Biophys Sin (Shanghai) 2009, 41:527–534.

34. Raaphorst FM: Self-renewal of hematopoietic and leukemic stem cells: a central role for the Polycomb-group gene Bmi-1. Trends Immunol 2003, 24:522–524.

35. Grinstein E, Mahotka C: Stem cell divisions controlled by the proto-oncogene BMI-1. J Stem Cells 2009, 4:141–146.

36. Weilemann A, Grau M, Erdmann T, Merkel O, Sobhiafshar U, Anagnostopoulos I, Hummel M, Siegert A, Hayford C, Madle H, et al: Essential role of IRF4 and MYC signaling for survival of anaplastic large cell lymphoma. Blood 2015, 125:124–132.

37. Kustikova OS, Geiger H, Li Z, Brugman MH, Chambers SM, Shaw CA, Pike-Overzet K, de Ridder D, Staal FJ, von Keudell G, et al: Retroviral vector insertion sites associated with dominant hematopoietic clones mark “stemness” pathways. Blood 2007, 109:1897–1907.

38. Deneault E, Cellot S, Faubert A, Laverdure JP, Frechette M, Chagraoui J, Mayotte N, Sauvageau M, Ting SB, Sauvageau G: A functional screen to identify novel effectors of hematopoietic stem cell activity. Cell 2009, 137:369–379.

39. Zhang H, Alberich-Jorda M, Amabile G, Yang H, Staber PB, Di Ruscio A, Welner RS, Ebralidze A, Zhang J, Levantini E, et al: Sox4 is a key oncogenic target in C/EBPalpha mutant acute myeloid leukemia. Cancer Cell 2013, 24:575–588.

40. Delmans M, Hemberg M: Discrete distributional differential expression (D3E)--a tool for gene expression analysis of single-cell RNA-seq data. BMC Bioinformatics 2016, 17:110.

41. Swords R, Freeman C, Giles F: Targeting the FMS-like tyrosine kinase 3 in acute myeloid leukemia. Leukemia 2012, 26:2176–2185.

42. Leung AY, Man CH, Kwong YL: FLT3 inhibition: a moving and evolving target in acute myeloid leukaemia. Leukemia 2013, 27:260–268.

43. Valk PJ, Verhaak RG, Beijen MA, Erpelinck CA, Barjesteh van Waalwijk van Doorn-Khosrovani S, Boer JM, Beverloo HB, Moorhouse MJ, van der Spek PJ, Lowenberg B, Delwel R: Prognostically useful gene-expression profiles in acute myeloid leukemia. N Engl J Med 2004, 350:1617–1628.

44. Cancer Genome Atlas Research N: Genomic and epigenomic landscapes of adult de novo acute myeloid leukemia. N Engl J Med 2013, 368:2059–2074.

45. Matsuo T, Kuriyama K, Miyazaki Y, Yoshida S, Tomonaga M, Emi N, Kobayashi T, Miyawaki S, Matsushima T, Shinagawa K, et al: The percentage of myeloperoxidase-positive blast cells is a strong independent prognostic factor in acute myeloid leukemia, even in the patients with normal karyotype. Leukemia 2003, 17:1538–1543.

46. Matsuo T, Cox C, Bennett JM: Prognostic significance of myeloperoxidase positivity of blast cells in acute myeloblastic leukemia without maturation (FAB: M1): an ECOG study. Hematol Pathol 1989, 3:153–158.

47. Itonaga H, Imanishi D, Wong YF, Sato S, Ando K, Sawayama Y, Sasaki D, Tsuruda K, Hasegawa H, Imaizumi Y, et al: Expression of myeloperoxidase in acute myeloid leukemia blasts mirrors the distinct DNA methylation pattern involving the downregulation of DNA methyltransferase DNMT3B. Leukemia 2014, 28:1459–1466.

48. Suic M, Boban D, Markovic-Glamocak M, Petrovecki M, Marusic M, Labar B: Prognostic significance of cytochemical analysis of leukemic M2 blasts. Med Oncol Tumor Pharmacother 1992, 9:41–45.

49. Levine JH, Simonds EF, Bendall SC, Davis KL, Amir el AD, Tadmor MD, Litvin O, Fienberg HG, Jager A, Zunder ER, et al: Data-Driven Phenotypic Dissection of AML Reveals Progenitor-like Cells that Correlate with Prognosis. Cell 2015, 162:184–197.

50. Schnerch D, Yalcintepe J, Schmidts A, Becker H, Follo M, Engelhardt M, Wasch R: Cell cycle control in acute myeloid leukemia. Am J Cancer Res 2012, 2:508–528.

51. Ye M, Zhang H, Amabile G, Yang H, Staber PB, Zhang P, Levantini E, Alberich-Jorda M, Zhang J, Kawasaki A, Tenen DG: C/EBPa controls acquisition and maintenance of adult haematopoietic stem cell quiescence. Nat Cell Biol 2013, 15:385–394.

52. Pabst T, Mueller BU, Zhang P, Radomska HS, Narravula S, Schnittger S, Behre G, Hiddemann W, Tenen DG: Dominant-negative mutations of CEBPA, encoding CCAAT/enhancer binding protein-alpha (C/EBPalpha), in acute myeloid leukemia. Nat Genet 2001, 27:263–270.

53. Tenen DG: Myeloid differentiation and the leukemia-initiating cell. Leuk Suppl 2014, 3:S25–26.

54. Okita K, Ichisaka T, Yamanaka S: Generation of germline-competent induced pluripotent stem cells. Nature 2007, 448:313–317.

55. Maddox J, Shakya A, South S, Shelton D, Andersen JN, Chidester S, Kang J, Gligorich KM, Jones DA, Spangrude GJ, et al: Transcription factor Oct1 is a somatic and cancer stem cell determinant. PLoS Genet 2012, 8:e1003048.

56. Tantin D: Oct transcription factors in development and stem cells: insights and mechanisms. Development 2013, 140:2857–2866.

57. Somerville TD, Wiseman DH, Spencer GJ, Huang X, Lynch JT, Leong HS, Williams EL, Cheesman E, Somervaille TC: Frequent Derepression of the Mesenchymal Transcription Factor Gene FOXC1 in Acute Myeloid Leukemia. Cancer Cell 2015, 28:329–342.

58. Cante-Barrett K, Pieters R, Meijerink JP: Myocyte enhancer factor 2C in hematopoiesis and leukemia. Oncogene 2014, 33:403–410.

59. Pon JR, Marra MA: MEF2 transcription factors: developmental regulators and emerging cancer genes. Oncotarget 2016, 7:2297–2312.

60. Bungartz G, Land H, Scadden DT, Emerson SG: NF-Y is necessary for hematopoietic stem cell proliferation and survival. Blood 2012, 119:1380–1389.

61. Gazit R, Garrison BS, Rao TN, Shay T, Costello J, Ericson J, Kim F, Collins JJ, Regev A, Wagers AJ, et al: Transcriptome analysis identifies regulators of hematopoietic stem and progenitor cells. Stem Cell Reports 2013, 1:266–280.

62. Shimamoto T, Ohyashiki JH, Ohyashiki K, Kawakubo K, Kimura N, Nakazawa S, Toyama K: GATA-1, GATA-2, and stem cell leukemia gene expression in acute myeloid leukemia. Leukemia 1994, 8:1176–1180.

63. North TE, de Bruijn MF, Stacy T, Talebian L, Lind E, Robin C, Binder M, Dzierzak E, Speck NA: Runx1 expression marks long-term repopulating hematopoietic stem cells in the midgestation mouse embryo. Immunity 2002, 16:661–672.

64. Tsuzuki S, Hong D, Gupta R, Matsuo K, Seto M, Enver T: Isoform-specific potentiation of stem and progenitor cell engraftment by AML1/RUNX1. PLoS Med 2007, 4:e172.

65. He X, Wang Q, Cen J, Qiu H, Sun A, Chen S, Wu D: Predictive value of high EVI1 expression in AML patients undergoing myeloablative allogeneic hematopoietic stem cell transplantation in first CR. Bone Marrow Transplant 2016.

66. Kataoka K, Sato T, Yoshimi A, Goyama S, Tsuruta T, Kobayashi H, Shimabe M, Arai S, Nakagawa M, Imai Y, et al: Evi1 is essential for hematopoietic stem cell self-renewal, and its expression marks hematopoietic cells with long-term multilineage repopulating activity. J Exp Med 2011, 208:2403–2416.

67. Renneville A, Roumier C, Biggio V, Nibourel O, Boissel N, Fenaux P, Preudhomme C: Cooperating gene mutations in acute myeloid leukemia: a review of the literature. Leukemia 2008, 22:915–931.

68. Zhang X, Lian Z, Padden C, Gerstein MB, Rozowsky J, Snyder M, Gingeras TR, Kapranov P, Weissman SM, Newburger PE: A myelopoiesis-associated regulatory intergenic noncoding RNA transcript within the human HOXA cluster. Blood 2009, 113:2526–2534.

69. Diaz-Beya M, Brunet S, Nomdedeu J, Pratcorona M, Cordeiro A, Gallardo D, Escoda L, Tormo M, Heras I, Ribera JM, et al: The lincRNA HOTAIRM1, located in the HOXA genomic region, is expressed in acute myeloid leukemia, impacts prognosis in patients in the intermediate-risk cytogenetic category, and is associated with a distinctive microRNA signature. Oncotarget 2015, 6:31613–31627.

70. Mayer IA, Arteaga CL: The PI3K/AKT Pathway as a Target for Cancer Treatment. Annu Rev Med 2016, 67:11–28.

71. Pollak M: Insulin and insulin-like growth factor signalling in neoplasia. Nat Rev Cancer 2008, 8:915–928.

72. Guo W, Schubbert S, Chen JY, Valamehr B, Mosessian S, Shi H, Dang NH, Garcia C, Theodoro MF, Varella-Garcia M, Wu H: Suppression of leukemia development caused by PTEN loss. Proc Natl Acad Sci U S A 2011, 108:1409–1414.

73. Borel C, Ferreira PG, Santoni F, Delaneau O, Fort A, Popadin KY, Garieri M, Falconnet E, Ribaux P, Guipponi M, et al: Biased allelic expression in human primary fibroblast single cells. Am J Hum Genet 2015, 96:70–80.

74. Langmead B, Salzberg SL: Fast gapped-read alignment with Bowtie 2. Nat Methods 2012, 9:357–359.

75. Kim D, Pertea G, Trapnell C, Pimentel H, Kelley R, Salzberg SL: TopHat2: accurate alignment of transcriptomes in the presence of insertions, deletions and gene fusions. Genome Biol 2013, 14:R36.

76. R Development Core Team: R: A Language and Environment for Statistical Computing. R Foundation for Statistical Computing, Vienna, Austria 2011.

77. Macosko EZ, Basu A, Satija R, Nemesh J, Shekhar K, Goldman M, Tirosh I, Bialas AR, Kamitaki N, Martersteck EM, et al: Highly Parallel Genome-wide Expression Profiling of Individual Cells Using Nanoliter Droplets. Cell 2015, 161:1202–1214.

78. Delmans M, Hemberg M: Discrete distributional differential expression (D(3)E) - a tool for gene expression analysis of single-cell RNA-seq data. BMC Bioinformatics 2016, 17:110.

79. Dar RD, Razooky BS, Singh A, Trimeloni TV, McCollum JM, Cox CD, Simpson ML, Weinberger LS: Transcriptional burst frequency and burst size are equally modulated across the human genome. Proc Natl Acad Sci U S A 2012, 109:17454–17459.

80. Dennis G, Jr., Sherman BT, Hosack DA, Yang J, Gao W, Lane HC, Lempicki RA: DAVID: Database for Annotation, Visualization, and Integrated Discovery. Genome Biol 2003, 4:P3.

81. Ben-Ari Fuchs S, Lieder I, Stelzer G, Mazor Y, Buzhor E, Kaplan S, Bogoch Y, Plaschkes I, Shitrit A, Rappaport N, et al: GeneAnalytics: An Integrative Gene Set Analysis Tool for Next Generation Sequencing, RNAseq and Microarray Data. OMICS 2016, 20:139–151.

82. Subramanian A, Kuehn H, Gould J, Tamayo P, Mesirov JP: GSEA-P: a desktop application for Gene Set Enrichment Analysis. Bioinformatics 2007, 23:3251–3253.

83. Franceschini A, Szklarczyk D, Frankild S, Kuhn M, Simonovic M, Roth A, Lin J, Minguez P, Bork P, von Mering C, Jensen LJ: STRING v9.1: protein-protein interaction networks, with increased coverage and integration. Nucleic Acids Res 2013, 41:D808–815.

84. Shannon P, Markiel A, Ozier O, Baliga NS, Wang JT, Ramage D, Amin N, Schwikowski B, Ideker T: Cytoscape: a software environment for integrated models of biomolecular interaction networks. Genome Res 2003, 13:2498–2504.

85. Huang da W, Sherman BT, Lempicki RA: Systematic and integrative analysis of large gene lists using DAVID bioinformatics resources. Nat Protoc 2009, 4:44–57.

86. Huang da W, Sherman BT, Lempicki RA: Bioinformatics enrichment tools: paths toward the comprehensive functional analysis of large gene lists. Nucleic Acids Res 2009, 37:1–13.

87. Cancer Genome Atlas Research N, Weinstein JN, Collisson EA, Mills GB, Shaw KR, Ozenberger BA, Ellrott K, Shmulevich I, Sander C, Stuart JM: The Cancer Genome Atlas Pan-Cancer analysis project. Nat Genet 2013, 45:1113–1120.

## SUPPLEMENTAL REFERENCES

Bonardi, F., Fusetti, F., Deelen, P., van Gosliga, D., Vellenga, E., and Schuringa, J.J. (2013). A proteomics and transcriptomics approach to identify leukemic stem cell (LSC) markers. Mol Cell Proteomics 12, 626–637.

Boquest, A.C., Shahdadfar, A., Fronsdal, K., Sigurjonsson, O., Tunheim, S.H., Collas, P., and Brinchmann, J.E. (2005). Isolation and transcription profiling of purified uncultured human stromal stem cells: alteration of gene expression after in vitro cell culture. Mol Biol Cell 16, 1131–1141.

Eppert, K., Takenaka, K., Lechman, E.R., Waldron, L., Nilsson, B., van Galen, P., Metzeler, K.H., Poeppl, A., Ling, V., Beyene, J., et al. (2011). Stem cell gene expression programs influence clinical outcome in human leukemia. Nat Med 17, 1086–1093.

Gal, H., Amariglio, N., Trakhtenbrot, L., Jacob-Hirsh, J., Margalit, O., Avigdor, A., Nagler, A., Tavor, S., Ein-Dor, L., Lapidot, T., et al. (2006). Gene expression profiles of AML derived stem cells; similarity to hematopoietic stem cells. Leukemia 20, 2147–2154.

Gentles, A.J., Plevritis, S.K., Majeti, R., and Alizadeh, A.A. (2010). Association of a leukemic stem cell gene expression signature with clinical outcomes in acute myeloid leukemia. JAMA 304, 2706–2715.

Georgantas, R.W., 3rd, Tanadve, V., Malehorn, M., Heimfeld, S., Chen, C., Carr, L., Martinez-Murillo, F., Riggins, G., Kowalski, J., and Civin, C.I. (2004). Microarray and serial analysis of gene expression analyses identify known and novel transcripts overexpressed in hematopoietic stem cells. Cancer Res 64, 4434–4441.

Jung, N., Dai, B., Gentles, A.J., Majeti, R., and Feinberg, A.P. (2015). An LSC epigenetic signature is largely mutation independent and implicates the HOXA cluster in AML pathogenesis. Nat Commun 6, 8489.

Levine, J.H., Simonds, E.F., Bendall, S.C., Davis, K.L., Amir el, A.D., Tadmor, M.D., Litvin, O., Fienberg, H.G., Jager, A., Zunder, E.R., et al. (2015). Data-Driven Phenotypic Dissection of AML Reveals Progenitor-like Cells that Correlate with Prognosis. Cell 162, 184–197.

Pelosi, E., Castelli, G., and Testa, U. (2015). Targeting LSCs through membrane antigens selectively or preferentially expressed on these cells. Blood Cells Mol Dis 55, 336–346.

Saito, Y., Kitamura, H., Hijikata, A., Tomizawa-Murasawa, M., Tanaka, S., Takagi, S., Uchida, N., Suzuki, N., Sone, A., Najima, Y., et al. (2010). Identification of therapeutic targets for quiescent, chemotherapy-resistant human leukemia stem cells. Sci Transl Med 2, 17ra19.

